# Agent based models to investigate cooperation between cancer cells

**DOI:** 10.1101/031070

**Authors:** Joao B. Xavier, William K. Chang

## Abstract

We present a type of agent-based model that uses off-lattice spheres to represent individual cells in a solid tumor. The model calculates chemical gradients and determines the dynamics of the tumor as emergent properties of the interactions between the cells. As an example, we present an investigation of cooperation among cancer cells where cooperators secrete a growth factor that is costly to synthesize. Simulations reveal that cooperation is favored when cancer cells from the same lineage stay in close proximity. The result supports the hypothesis that kin selection, a theory that explains the evolution of cooperation in animals, also applies to cancers.

## I. Cooperation among cancer cells

It has long been hypothesized that tumors represent complex societies of mutualistically-interacting cell types[1, 2]. Recent research shows that many of the malignant phenotypes of tumours known as the ‘hallmarks of cancer’[3, 4] represent population-level behaviors. Experiments reveal that tumor growth may be driven by a subpopulation of tumor cells[5] and that tumorigenesis may require the participation of distinct phenotypic subpopulations[6]. Such interactions constitute a potential mechanism for clonal interference and provide an explanation for experimentally observed intratumoral heterogeneity[7, 8].

In addition to heterogeneity within the cancer cell population giving rise to synergistic interactions, the tumor microenvironment is populated by non-transformed host tissue and immune cells, which play important roles in determining tumor progression[9, 10] and treatment response[11]. Feedback in tumor-microenvironment interactions leads to the co-evolution of cancer cells and their environment[12]. As with many dynamical systems involving feedback interactions, the output of tumor-microenvironment interactions can potentially be chaotic and unpredictable, as can consequences of attempts to perturb the system with conventional or targeted treatments.

Given the complex multi-scale organization of the tumor ecosystem, mathematical modeling can be a powerful tool to understand how the dynamics of the cellular ecosystem drive tumor progression and the consequences of perturbing this system with traditional or targeted therapy. In particular, applying mathematical frameworks derived from ecology and evolutionary theory allows the cancer biologist to leverage existing methods to analyze the collective behavior, population dynamics, and evolutionary dynamics of cellular systems in a way that is robust with respect to the constantly shifting genetic and molecular background within and between tumors[8, 13].

## II. An application of agent-based models

Individual cells within each subpopulation may experience different microenvironmental conditions or stochastic fluctuations, and consequently display different phenotypes and behaviors. In this case, the population output integrates, through interactions between individuals, these individually noisy stimuli-response calculations.

Many models simulating the population level behavior as emergent from varying individual cell activities fall under the general category of agent-based models (ABM)[14]. These models, constituting a ‘bottom up’ simulation approach, have an additional advantage of being integrable with a continuum representation of diffusible chemical gradients, for example of cytokines or glucose. Such integrated models are sometimes referred to as hybrid discrete-differential (HDD) models[15]. Between the discrete-space representation of cells and their spatial organization, and the continuum representation of chemical species, HDD models have the potential to simulate the aggregate output of a multitude of regulatory mechanisms and interactions.

ABMs are useful for exploring the effect of spatial structure on population and evolutionary dynamics. In microbial and tissue systems, diffusive substances such as nutrients and growth factors often have a decisive effect on population dynamics. The spatial distribution of resources and individuals thus determine the dynamics of the ecosystem, possibly in ways that contradict the predictions of mean-field models. For instance, spatial structure can explain the emergence and maintenance of public-goods cooperative behavior in evolving populations if cooperative individuals aggregate and the public good is spatially limited[16]. This is readily demonstrated in agent-based spatial simulations for the example of bacterial colonies[17].

Using a variant of this bacterial model, it is also possible to show the emergence of a cooperative subclone in a two-dimensional spherical tumour (Fig. 1). Conversely, the emergence of spatial structure can be evidence of interspecific interactions [18]. The presence of intricate spatial structure and nonuniform distribution of cell types within tumors[19, 20] may point to the dominance of interspecific effects within tumor cell populations.

**Figure 1.**
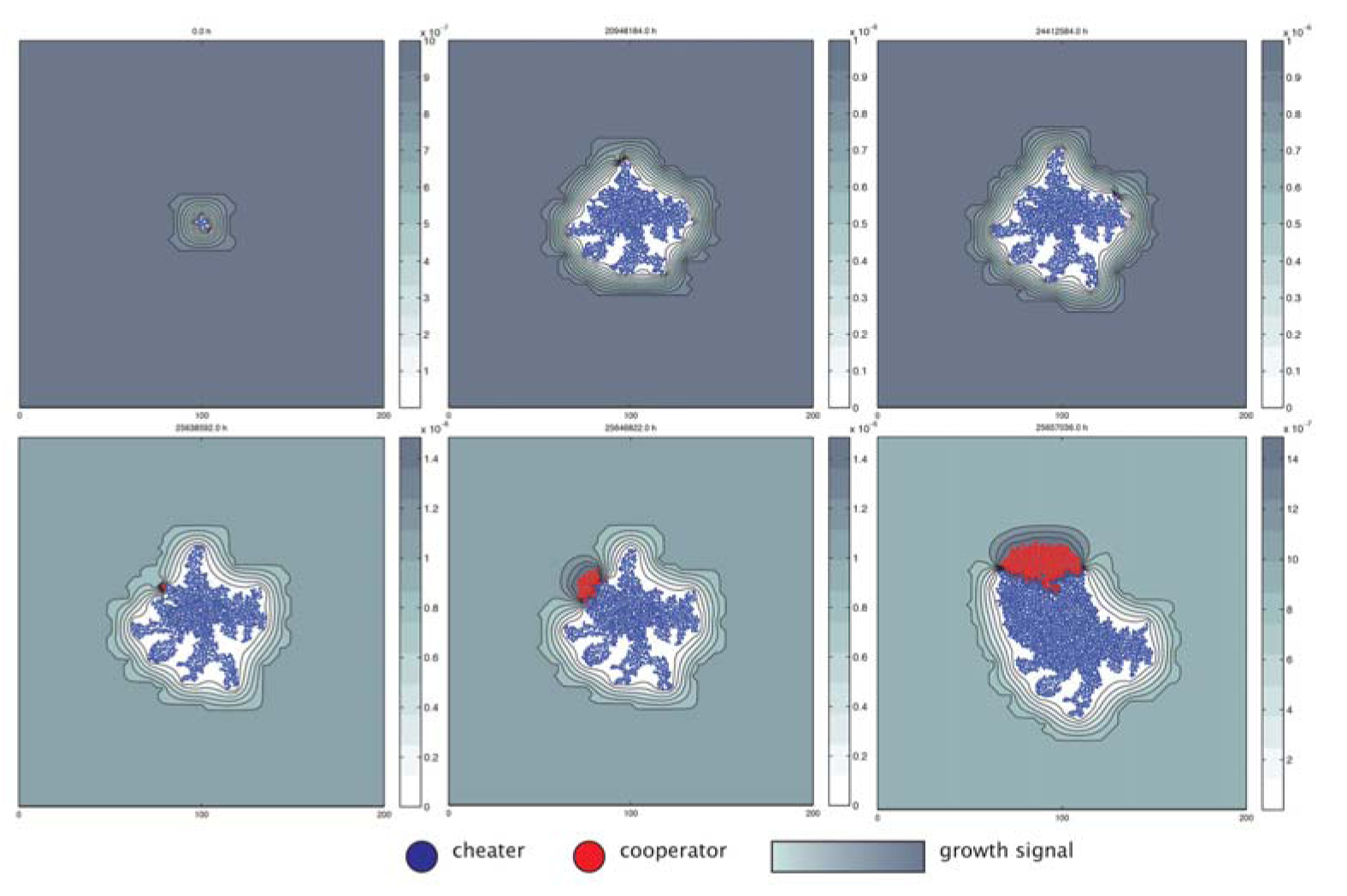
Agent based models reveal that cooperation between cancer cells can persist if cooperators are in close proximity so that they share cooperative secretions with each other. The simulation is initiated with one non-cooperative (‘cheater’) tumor cell at the center of the simulation space. Growth signal is initially uniformly distributed throughout the space, but gradients emerge as the signal is consumed by growing cells. Stochastic emergence of a cooperative subtype, which expresses the growth signal at a cost to its own growth, at the edge of the expanding tumor leads to a locally high concentration of growth signal and overall acceleration of cell growth. The emergent spatial structure—aggregation of cooperators and consequent segregation of public goods—stabilizes the cooperative subpopulation.

## III. Methods

For the simulations shown in Fig 1. we used an ABM where every cell is represented by an independent agent (a ‘virtual cell’) with a defined size and location represented in spatial continuous coordinates. The behavior of each agent is determined by a set of rules that mimics the behavior of real cells: cells grow, divide, move due to mechanical displacement and forces applied by other cells, consume nutrients and secrete signaling molecules and growth factors that enhance growth of neighboring cells. Importantly, the kinetic rates for each of the processes are determined by the local microenvironment that each cell experiences.

The microenvironment, i.e. the local concentrations of each solute such as nutrients and growth factors, is determined by solving a system of coupled partial-differential equations (PDEs) that take into account diffusion and reaction (i.e. production and consumption) of each solute.

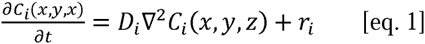

In equation 1, C_i_(x,y,z) represents the concentration of solute i at location (x, y, z) in space, D_i_ is the diffusivity coefficient of that solute and ri is the net biochemical reaction rate for that solute (r_i_>0 if solute ‘i’ is being produced, which can be the case for growth factors, and r_i_<0 if solute ‘i’ is being consumed, as in the case of nutrients). The solute species can be any relevant chemical species such as oxygen and glucose that are consumed by the cells, but also substances that are produced by the cells such as growth factors or metabolic waste products such as lactate. The framework allows any number of solute substances to be included.

### Other applications for this model

The true power of agent-based models comes from their generality. The particular framework described here was first developed to model bacterial communities[21]. It has been applied to a range of systems, from modeling wastewater treatment reactors[22], to investigating mechanisms of biofilm dispersal[23] to the study of bacterial cooperation[24]. More recently, these agent-based models have been applied to investigate tumor-stromal interactions and in particular the emergence of spatial structure in tumor-associated macrophages[18].

## References (Highly Relevant up to 1 page)

[1] G. H. Heppner, “Tumor cell societies,” Journal of the National Cancer Institute, vol. 81, pp. 648–649, 1989.

[2] R. Axelrod, D. E. Axelrod, and K. J. Pienta, “Evolution of cooperation among tumor cells,” Proceedings of the National Academy of Sciences, vol. 103, pp. 13474–13479, 2006.

[3] D. Hanahan and R. A. Weinberg, “The hallmarks of cancer,” cell, vol. 100, pp. 57–70, 2000.

[4] D. Hanahan and R. A. Weinberg, “Hallmarks of cancer: the next generation,” cell, vol. 144, pp. 646–674, 2011.

[5] A. Marusyk, D. P. Tabassum, P. M. Altrock, V. Almendro, F. Michor, and K. Polyak, “Non-cell-autonomous driving of tumour growth supports sub-clonal heterogeneity,” Nature, vol. 514, pp. 54–58, 2014.

[6] A. S. Cleary, T. L. Leonard, S. A. Gestl, and E. J. Gunther, “Tumour cell heterogeneity maintained by cooperating subclones in Wnt-driven mammary cancers,” Nature, vol. 508, pp. 113–117, 2014.

[7] E. C. de Bruin, N. McGranahan, R. Mitter, M. Salm, D. C. Wedge, L. Yates, et al., “Spatial and temporal diversity in genomic instability processes defines lung cancer evolution,” Science, vol. 346, pp. 251–256, 2014.

[8] K. S. Korolev, J. B. Xavier, and J. Gore, “Turning ecology and evolution against cancer,” NatRev Cancer, vol. 14, pp. 371–80, May 2014.

[9] D. Hanahan and L. M. Coussens, “Accessories to the crime: functions of cells recruited to the tumor microenvironment,” Cancer cell, vol. 21, pp. 309–322, 2012.

[10] D. F. Quail and J. A. Joyce, “Microenvironmental regulation of tumor progression and metastasis,” Nature medicine, vol. 19, pp. 1423–1437, 2013.

[11] D. G. DeNardo, D. J. Brennan, E. Rexhepaj, B. Ruffell, S. L. Shiao, S. F. Madden, et al., “Leukocyte complexity predicts breast cancer survival and functionally regulates response to chemotherapy,” Cancer discovery, vol. 1, pp. 54–67, 2011.

[12] K. Polyak, I. Haviv, and I. G. Campbell, “Co-evolution of tumor cells and their microenvironment,” Trends in Genetics, vol. 25, pp. 30–38, 2009.

[13] C. Swanton, “Intratumor heterogeneity: evolution through space and time,” Cancer research, vol. 72, pp. 4875–4882, 2012.

[14] L. Zhang, Z. Wang, J. A. Sagotsky, and T. S. Deisboeck, “Multiscale agent-based cancer modeling,” Journal of mathematical biology, vol. 58, pp. 545–559, 2009.

[15] A. R. Anderson, “A hybrid mathematical model of solid tumour invasion: the importance of cell adhesion,” Mathematical Medicine and Biology, vol. 22, pp. 163–186, 2005.

[16] J. Y. Wakano, M. A. Nowak, and C. Hauert, “Spatial dynamics of ecological public goods,” Proceedings of the National Academy of Sciences, vol. 106, pp. 7910–7914, 2009.

[17] C. D. Nadell, K. R. Foster, and J. B. Xavier, “Emergence of spatial structure in cell groups and the evolution of cooperation,” PLoS Comput Biol, vol. 6, p. e1000716, Mar 2010.

[18] C. Carmona-Fontaine, V. Bucci, L. Akkari, M. Deforet, J. A. Joyce, and J. B. Xavier, “Emergence of spatial structure in the tumor microenvironment due to the Warburg effect,” Proceedings of the National Academy of Sciences, vol. 110, pp. 19402–19407, 2013.

[19] V. Gocheva, H.-W. Wang, B. B. Gadea, T. Shree, K. E. Hunter, A. L. Garfall, et al., “IL-4 induces cathepsin protease activity in tumor-associated macrophages to promote cancer growth and invasion,” Genes & development, vol. 24, pp. 241–255, 2010.

[20] J. Zhang, J. Fujimoto, J. Zhang, D. C. Wedge, X. Song, J. Zhang, et al., “Intratumor heterogeneity in localized lung adenocarcinomas delineated by multiregion sequencing,” Science, vol. 346, pp. 256–259, 2014.

[21] J. B. Xavier, C. Picioreanu, and M. C. van Loosdrecht, “A framework for multidimensional modelling of activity and structure of multispecies biofilms,” Environ Microbiol, vol. 7, pp. 1085–103, Aug 2005.

[22] J. B. Xavier, M. K. De Kreuk, C. Picioreanu, and M. C. Van Loosdrecht, “Multi-scale individual-based model of microbial and bioconversion dynamics in aerobic granular sludge,” Environmental science & technology, vol. 41, pp. 6410–6417, 2007.

[23] J. B. Xavier, C. Picioreanu, S. A. Rani, M. C. van Loosdrecht, and P. S. Stewart, “Biofilm-control strategies based on enzymic disruption of the extracellular polymeric substance matrix--a modelling study,” Microbiology, vol. 151, pp. 3817–32, Dec 2005.

[24] J. B. Xavier and K. R. Foster, “Cooperation and conflict in microbial biofilms,” Proc Natl Acad Sci USA, vol. 104, pp. 876–81, Jan 16 2007.

